# Extracorporeal Placental Support in a Sheep’s Gravid Uterus

**DOI:** 10.1101/2025.07.02.662865

**Authors:** Jose H Salazar, Jennifer M Schuh, Araceli Morelos, Ivonne Monroy, Emmanuel Abebrese, Natalie Stocke, Nicholas Starkey, Deron Jones, G Ganesh Konduri

## Abstract

**BACKGROUND:** The placenta is a vital organ in animal development, yet its function is understudied. Existing ex vivo placental support models study the organ in isolation, without the uterus and fetus. We report our early experience with a novel model of extracorporeal placental support by perfusing a sheep’s gravid uterus.

**METHODS:** A sheep gravid hysterectomy was performed after cannulating the uterine vessels. The isolated uterus was maintained in a warm crystalloid tank. Oxygenation, electrolyte supplementation, and glucose were provided through an extracorporeal life support circuit, while monitoring the fetal heart rate and pressure. Our primary objective was to establish the feasibility of the model, confirm the ability to transition the organ to ex vivo support (fetal survival for >30 minutes after hysterectomy), and measure the duration of fetal survival after uterine explantation.

**RESULTS:** A total of 51 surgeries were performed, and 76% were successfully transitioned to extracorporeal support. Growing experience throughout the study period led to technical adaptations and more prolonged survival. Uteri with a singleton fetus had longer mean survival times, when compared to twins (253 minutes vs. 104 minutes, p=.019). Anemia and coagulopathy were frequently encountered, and most of the uteri had significant clot formation around the placentomes at the end of the experiment.

**CONCLUSION:** Using a model of extracorporeal placental support in sheep, we were able to maintain a live fetus within the uterus for more than 16 hours after hysterectomy. This technique is feasible and could provide an avenue to study fetoplacental physiology in an ex vivo environment.

## BACKGROUND

The placenta is a crucial organ for animal development. Despite its relevance, there are disproportionately fewer investigations of its function when compared to other solid organs. The ultimate marker of placental function is a healthy fetus, and in- or ex-vivo placental studies are limited by challenges in accessing the gravid uterus and ethical considerations. The sheep has been used as an alternative model in exploring fetoplacental physiology. Several commonalities with humans include fetal size, litter numbers, cord structure, pulmonary development, and fetal circulation.

A model of ex vivo placental support in sheep does not have direct translation to human use, but the insight that it can provide into fetoplacental physiology can lead into multiple research avenues which may help better understand placental function and its behavior when supported extracorporeally. In this manuscript, we seek to describe our experience with extracorporeal placental support by perfusing a sheep’s gravid uterus. We discuss the technical challenges, hemodynamic changes, fetal survival, and knowledge obtained from this early experience.

## METHODS

### Animal model

The extracorporeal placental support (EPS) model is based on sheep for several reasons, some of these are: litter number, uterine and fetal size, vessel diameter, and extensive reports in the literature on pregnant sheep. Cheviot, Suffolk, and Dorset breeds were used during the experiments. Surgery was conducted in singleton and twin pregnancies between 89 and 120 days of gestation to match the timing of the transition between the canalicular and saccular stages of lung development, which corresponds to 22 to 26 weeks of gestation in humans.(1) Animals were fasted for 12 hours prior to the surgery, and an intramuscular injection of tiletamine and zolazepam was delivered. Propofol was then administered, and anesthesia was maintained with isoflurane. 100 mg of indomethacin was given per rectum before laparotomy to prevent uterine contractions. The study was approved by the Institutional Animal Care and Use Committee (AUA00007678) at the Medical College of Wisconsin.

### Surgical approach

A low transverse laparotomy was performed and the uterus was externalized through the wound. A hysterotomy was then made, and monitoring lines were placed to obtain real-time measurements of fetal heart rate. A pressure catheter (2fr, Millar, Houston) was placed in a placentomic artery or, on occasion, in the fetal carotid artery. The hysterotomy was closed, and the bilateral uterine vessels were dissected and cannulated to transition the gravid uterus to extracorporeal support. Heparin (80u/kg) was given to the ewe prior to cannulation. The hysterectomy was completed by transecting the cervix, and the uterus was transferred to a warm crystalloid tank to continue support and control the temperature.**(Figure 1)** The ewe was bled to maintain the extracorporeal circuit with sheep’s whole blood and then euthanized with a pentobarbital-containing euthanasia solution.

**Figure 1.**
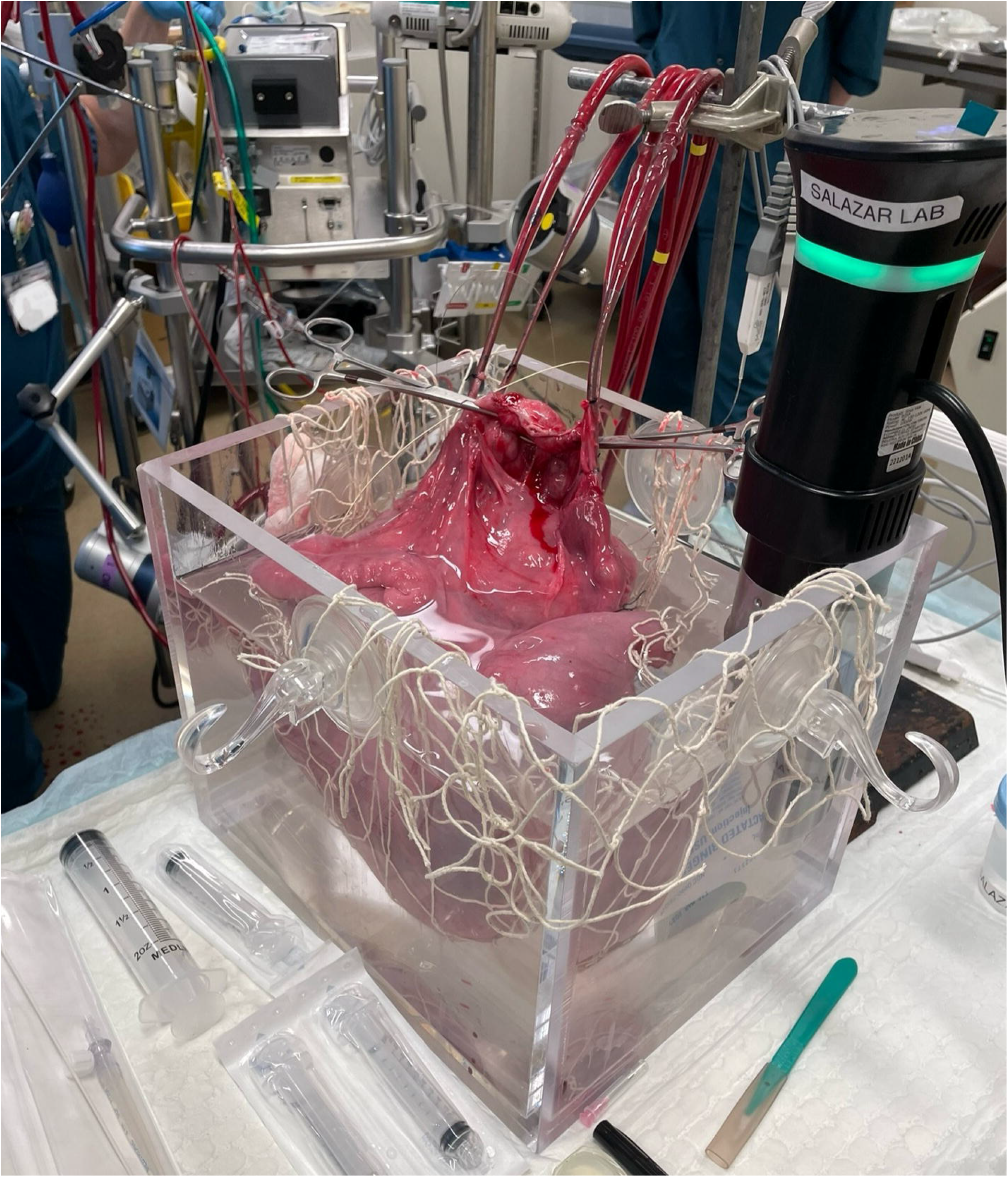
Gravid uterus under extracorporeal placental support

### Extracorporeal Circuit

The circuit was composed of a centrifugal pump with an RF-32 pump head (Getinge, Stockholm), a pediatric Capiox RX25 oxygenator (Terumo, Ann Arbor) primed with 1 liter of crystalloid and 800mL of sheep whole blood. A Hemocor HPH hemoconcentrator (Terumo, Ann Arbor) was used to filter crystalloid and plasma and maintain an adequate hematocrit. A Hemotherm heater/cooler (Gentherm, Northville) maintained normothermia. CO2 was kept at normal ranges with sweep gas powered by an air and oxygen blender system (Sechrist, Anaheim). Flow through the native uterine arteries was checked before cannulation using PSB flow probes (Transonic, Ithaca), and venoarterial support was provided at a flow rate that reflects physiologic uterine flow (200-400 mL/min per artery) and titrated based on gas exchange.

### Uteroplacental support

Flow through the circuit was titrated by controlling the revolutions on the pump or adding fluid (i.e. sheep whole blood or crystalloid). We attempted to keep the hemoglobin level between 10 and 14 g/dL by hemoconcentrating the circuit. Serial blood samples were obtained to check electrolytes, activated clotting times (ACT), and blood gases using a Vetscan i-STAT (Abbott, Lake County). Calcium chloride or gluconate, sodium bicarbonate, and glucose were administered throughout the experiment. Heparin (50-500 units) was given if the ACT fell below 150 seconds. Tocolytics (i.e. magnesium sulfate or nicardipine) were infused to stop uterine contractions. Real-time measurements of fetal heart rate and circuit flow were monitored using PowerLab and displayed with LabChart (AdInstruments, NSW Australia).

### Outcomes

During this early experience, we sought to confirm the model’s feasibility and analyze the frequency of successful transitions of a gravid uterus to extracorporeal support. A successful transition was defined as a live fetus for >30 minutes after hysterectomy. On those that were successfully transitioned, we investigated for how long we could maintain adequate placental function, defined by fetal survival. The experiment was terminated when the fetal heart stopped, as confirmed by the absence of a readable heart rate by the pressure catheter and lack of cardiac activity on ultrasound. As the circuit is primed with crystalloid and whole blood, we were interested in the occurrence of anemia, defined as hemoglobin concentration <8g/dL.(2) There was no formal necropsy on the biological specimens.

## RESULTS

A total of 51 surgeries were performed to attempt extracorporeal support of a gravid uterus. (**Table 1)** Of these, 76% (n=39) were successfully transitioned to extracorporeal support (defined as a uterus with a live fetus >30 minutes after hysterectomy). Fetal survival after successfully transitioning the uterus ranged from 36 min to 980 min.

**TABLE 1.** Gestation details, hemoglobin (Hb), activated clotting time (ACT), and duration of each surgery.

| 1st tertile |  |  |  |  |  | 2nd tertile |  |  |  |  |  | 3rd tertile |  |  |  |  |  |
| --- | --- | --- | --- | --- | --- | --- | --- | --- | --- | --- | --- | --- | --- | --- | --- | --- | --- |
| n | Gestational age (days) | # of fetuses | Hb range (g/dL) | ACT range (seconds) | Time from explantation to fetal demise (minutes) | n | Gestational age (days) | # of fetuses | Hb range (g/dL) | ACT range (seconds) | Time from explantation to fetal demise (minutes) | n | Gestational age (days) | # of fetuses | Hb range (g/dL) | ACT range (seconds) | Time from explantation to fetal demise (minutes) |
| 1 | 115 | 2 | 6.1-7.8 | NA | 43 | 18 | 113 | 1 | NA | NA | 23 | 35 | 117 | 2 | 6.5-12.6 | 147-244 | 287 |
| 2 | 113 | 2 | 10.5 | NA | 47 | 19 | 117 | 2 | NA | 173 | 161 | 36 | 114 | 1 | NA | NA | - |
| 3 | 112 | 1 | NA | NA | 54 | 20 | 100 | 2 | NA | NA | 8 | 37 | 114 | 1 | 8.8-10.5 | >1000 | 126 |
| 4 | 100 | 2 | NA | NA | 42 | 21 | 103 | 2 | NA | NA | 41 | 38 | 114 | 1 | 9.2-13.6 | 139-178 | 297 |
| 5 | 107 | 2 | 7.1 | 101->1000 | 160 | 22 | 107 | 1 | <5-12.2 | 420 | 191 | 39 | 114 | 1 | 7.5 | >800 | 52 |
| 6 | 104 | 1 | <5 | NA | - | 23 | 110 | 1 | 5.4-12.2 | 420 | 156 | 40 | 113 | 1 | 7.1 | 104 | 45 |
| 7 | 94 | 1 | <5-<5 | 216 | 58 | 24 | 117 | 2 | NA | NA | 20 | 41 | 114 | 1 | 5.8-9.9 | 441->1000 | 181 |
| 8 | 89 | 1 | NA | NA | 70 | 25 | 102 | 1 | 5.8-13.6 | 160-820 | 404 | 42 | 113 | 1 | NA | NA | - |
| 9 | 115 | 2 | NA | NA | 38 | 26 | 106 | 2 | NA | NA | 79 | 43 | 111 | 1 | NA | NA | 19 |
| 10 | 109 | 2 | NA | NA | - | 27 | 116 | 2 | NA | NA | 25 | 44 | 113 | 1 | 11.2-14.2 | 147-170 | 196 |
| 11 | 113 | 2 | NA | NA | - | 28 | 100 | 1 | NA | NA | 18 | 45 | 113 | 1 | 7.5-11.9 | NA | 261 |
| 12 | 113 | 1 | <5-<5 | 156 | 89 | 29 | 105 | 1 | 7.5-7.8 | 151-1050 | 173 | 46 | 114 | 1 | 7.5-12.9 | 100-255 | 980 |
| 13 | 106 | 1 | 5.8 | 552 | 90 | 30 | 99 | 1 | 6.8-13.6 | 93-426 | 840 | 47 | 110 | 1 | 9.5 | 251 | 96 |
| 14 | 109 | 1 | NA | NA | 40 | 31 | 100 | 1 | 5.4-16.0 | 130-871 | 394 | 48 | 111 | 1 | 7-14.3 | 127-546 | 860 |
| 15 | 112 | 2 | 7.5-10.2 | 143 | 206 | 32 | 114 | 1 | NA | NA | 28 | 49 | 110 | 1 | NA | NA | 58 |
| 16 | 115 | 2 | 6.8 | 156 | 36 | 33 | 117 | 1 | 6.5-8.5 | NA | 128 | 50 | 113 | 1 | 10.5-10.9 | 147 | 131 |
| 17 | 110 | 1 | NA | NA | 43 | 34 | 114 | 1 | 8.5-9.5 | 131->700 | 115 | 51 | 114 | 1 | 6.1-13.6 | 178-364 | 955 |

Singleton pregnancies were transitioned to extracorporeal in 28 of the 35 (80%) singleton experiments, while twin pregnancies were transitioned in 11 of the 16 (69%) twin surgeries. Transitioned uteri with a singleton had longer fetal survival than twin pregnancies (n=11) with a mean duration in minutes of 253 vs. 104 (p=.019), respectively. Hemoglobin levels were tested in 29 experiments, and circuit anemia was a frequent issue, with 76% (n=22) of the uteri with a hemoglobin level <8g/dL at some point during ex vivo support.(2) Once we started using a hemoconcentrator, we were able to treat anemia more efficiently. 33% (n=13) of the experiments had ACT values >300 seconds. Electrolytes such as sodium and potassium were generally stable. Due to the type of i-STAT cartridges used, chloride was not frequently checked, but it was normal on the few occasions that we tested for it. Calcium was supplemented in most experiments, as blood used for the circuit was maintained in ethylenediaminetetraacetic acid (EDTA) bags, likely contributing to the hypocalcemia. Acidosis was common and managed with sodium bicarbonate infusions. Once lactate began rising, it was difficult to reverse the trend, as the physiology of the ex vivo model does not offer good hepatic or renal clearance.

The length of ex vivo placental support increased as the learning curve of this novel procedure advanced and technical adaptations were implemented. **(Figure 2)** The mean survival of successfully transitioned gravid uteri to extracorporeal circulation was 73 (SD 51), 244 (SD 229), and 323 (SD 341) (p=0.028) for each tertile, respectively.

**Figure 2.**
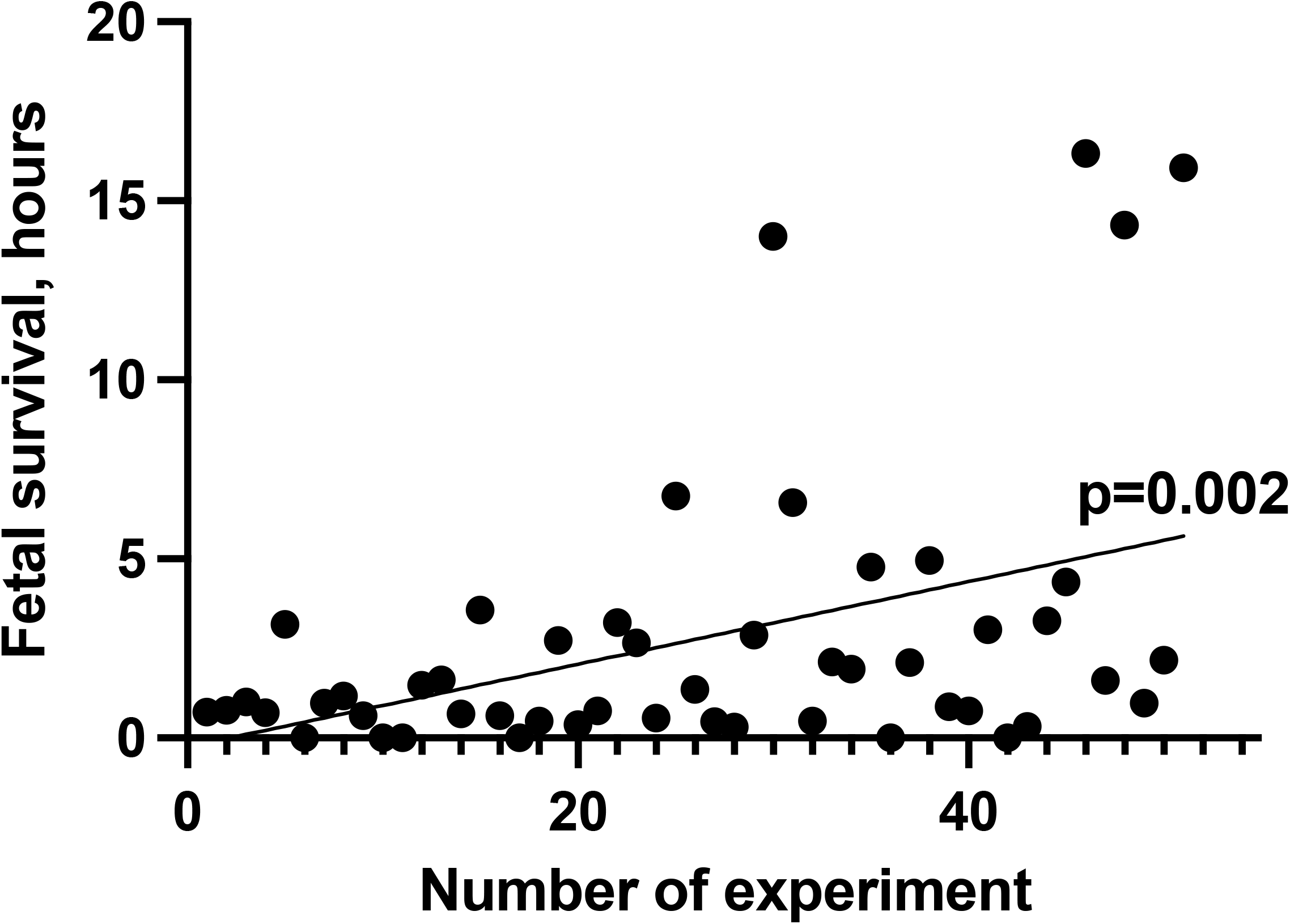
Fetal survival in each experiment. Line represents the linear regression slope.

While there was not a formal pathologic examination of each uterus and fetus, we believe that the causes of fetal demise at the beginning of our experience were mostly related to technical complications of cannulation (e.g. bleeding, poor inflow from cannula position), excessive uterine manipulation causing fetal distress, poor oxygen delivery from circuit anemia, and rapid fluctuations in glucose and electrolytes. We changed the design of the experiments throughout the experience to attempt to control for these causes of death and prolong survival. Some of these surgical and management adaptations were: hemoconcentration to avoid anemia, early management of contractions, microdosing heparin (<20 units per dose) to avoid circuit coagulopathy, increased frequency of electrolyte and gas exchange measurement, use of drips for electrolyte and glucose supplementation instead of bolus and utilizing singleton instead of twin pregnancies to facilitate uterine manipulation and retraction. During the longest experiments, we noticed a significant amount of clot formation around the placentomes, despite better-regulated ACT levels.

The hemodynamic course and laboratory values of the experiment that lasted the longest (980 minutes) are depicted in **Figures 3 and 4.**

**Figure 3.**
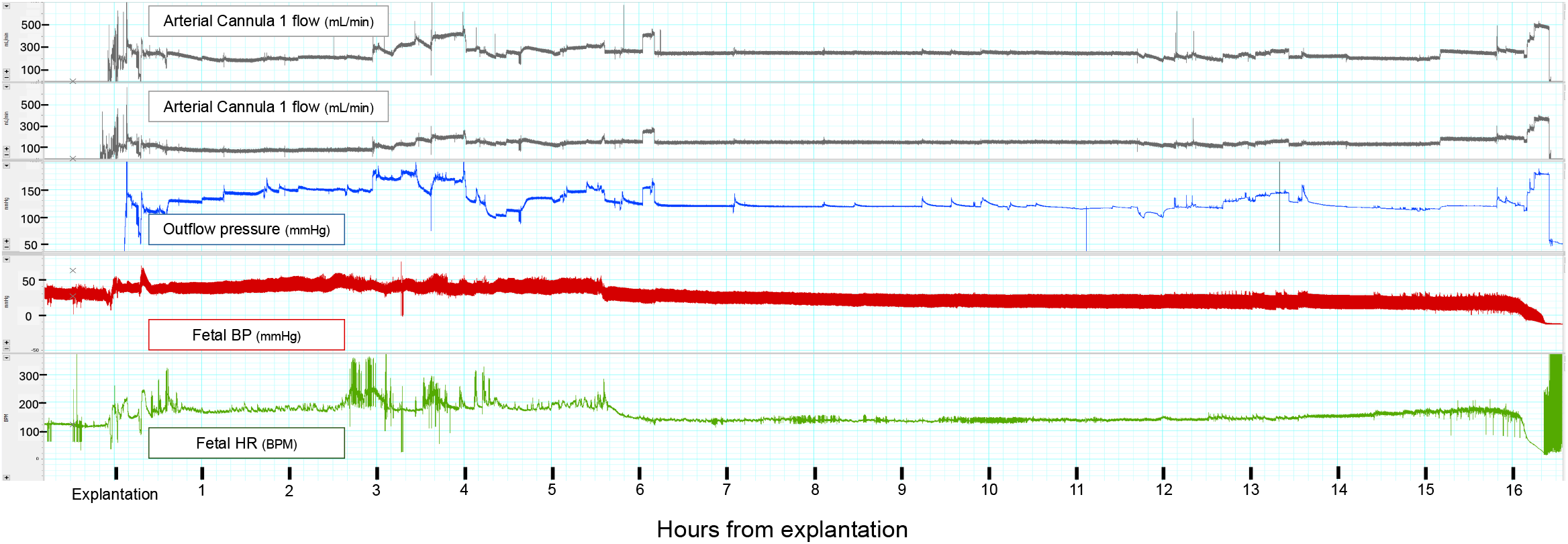
Hemodynamic monitoring of the extracorporeal placental support circuit and the fetus during a 16-hour experiment. BP, blood pressure; HR, heart rate.

**Figure 4.**
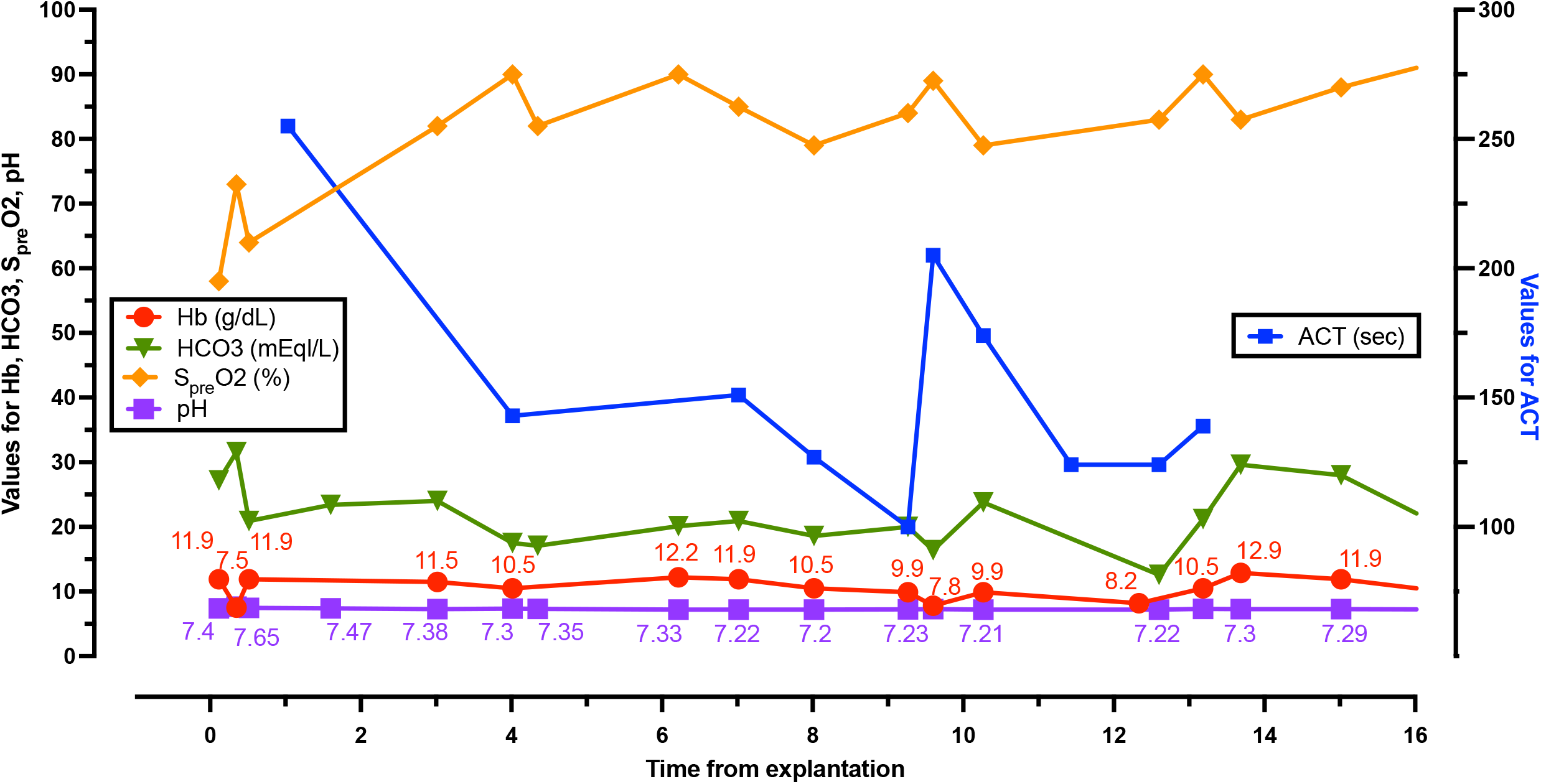
Laboratory values during a 16-hour experiment. Hb, hemoglobin; HCO3, bicarbonate; S_pre_O2, pre-oxygenator oxygen saturation; ACT, activated clotting time.

## DISCUSSION

Isolated organ perfusion has been well studied in solid organs, but there are very limited data on the tolerance of a gravid uterus to extracorporeal support. The term “ex vivo placental support” is commonly used to describe sophisticated models where one or several cotyledons are cannulated in a fresh placenta. This is typically used for experiments on solute filtration and less commonly for vascular or tissue injury studies.(3) A crucial indicator of placental function is fetal health, and such models do not include the fetus. With the EPS model, we perfuse a gravid uterus and study fetoplacental physiology without isolating the placenta or the fetus. We hereby report our early experience with this feasible model and describe some of the challenges and results of our experiments.

The best biomarker of placental function, in the setting of EPS, is fetal survival. We did not perform necropsies during this initial experience, but we hypothesize that the most common causes of early (within 1-2 hours) fetal demise were failure to recover from the hemodynamic instability associated with the manipulation, cannulation, explantation, acute electrolyte imbalances, flow disturbances, and anemia. While late (>2 hours) fetal demise was likely due to an inflammatory response from exposure to the circuit, placental failure from clot formation, or a humoral factor that we have yet to determine. Infection could be an important factor, but this is more likely to affect our model once more prolonged survivals are obtained. We performed intraoperative ultrasound in some of the experiments and noticed that fetal circulation was maintained, therefore, we do not believe that ductus arteriosus closure played a role. Now that we are achieving fetal survival for more than 12 hours, we plan to ascertain more physiologic variables and chemokine values that will offer a better understanding of the mechanisms of fetal death.

Some factors that we recognized during our experience that will inform future iterations of the model were: (1) the uterine vessels were large enough to provide the same flows that the uterus had in its native state (200-400 mL/min per artery), (2) we frequently noticed the presence of clot around the placentomes, but there was no evidence of bleeding in the fetus at the end of the experiments, suggesting adequate heparin filtration by the placenta, (3) while the sheep uterus is thought to be resistant to contractions during manipulation, we frequently noticed contractions, which were controlled with magnesium sulfate and nicardipine, (4) we were not able to revert acidosis in most cases, (5) a monitoring line in the placentomic arteries can provide access to fetal pressure and heart rate, but it is very sensitive to motion and position., and (6) even small doses of heparin (i.e. 50 units) in the circuit can significantly increase the ACT while on extracorporeal support.

A human gravid hysterectomy is a clinical extreme; therefore, the main limitation of our model is that it is not directly translational. Nevertheless, there are multiple applications where an ex vivo model of placental support can be utilized to improve current knowledge. Some of these include hypoxia/hyperoxia responses, anemia/hyperviscosity tolerance, clot formation and thrombogenesis, validation of imaging algorithms of flow and pulsatility, presence of placental biomarkers, and extrapolating findings of uterine support to solid organ preservation. Regarding clinical translation, several sophisticated models, where a sheep fetus receives extracorporeal support by cannulating umbilical and/or central vessels, are described with the term artificial womb technology (AWT).

AWT models, such as the Extra-uterine environment for neonatal development (EXTEND), the Ex-vivo uterine environment (EVE), and venovenous ECLS, are close to human use.(4–6) The data obtained from the EPS model may offer some insight into the interaction between an extracorporeal circuit and the fetoplacental unit, potentially offering data that could optimize AWT models.

The EPS model combines the challenges encountered in normothermic organ preservation and AWT. The EXTEND model of AWT has shown fetal survival for up to 4 weeks, but isolated organ perfusion has shorter perfusion times. Normothermic perfusion of human kidneys has been reported for up to 73 hours, human liver for 17 days, and sheep’s heart for 72 hours.(7–9) Different than perfusing the whole fetus, as is done in AWT, single-organ perfusion can lead to metabolic waste buildup, unregulated inflammatory activation, greater risk of hemolysis, lower capacity to repair tissue injury, and infection.(10–12) As we persist in our efforts to prolong placental support, we need to continue incorporating existing data on isolated organ perfusion and AWT. We believe that the EPS model can be optimized to achieve the longevity seen in isolated organ perfusion models, and we are in the process of understanding whether survival can be prolonged to lengths that are seen in AWT models.

The placenta is an understudied organ, and improving our knowledge of its function can translate into better outcomes for mothers and newborns. Using a model of extracorporeal placental support, we were able to maintain a live fetus within the uterus for more than 16 hours after explantation. This technique can provide an avenue to study fetoplacental physiology in an ex vivo environment. As experience with the model improves, fetal survival is prolonged, and the EPS model will open an opportunity for multiple avenues in placental research.

## Author contributions

JHS: Contributed to the concept/design, data analysis/interpretation, drafting the article, statistics, and data collection.

JMS: Contributed to the concept/design, data analysis/interpretation, drafting the article, and data collection.

AM: Contributed to data analysis/interpretation and data collection.

IM: Contributed to data analysis/interpretation, and data collection.

EA: Contributed to the data analysis/interpretation and data collection.

NS: Contributed with concept/design, and data collection.

NS: Contributed with concept/design, and data collection.

DJ: Contributed with concept/design, data analysis/interpretation, and data collection.

GGK: Contributed to the concept/design, data analysis/interpretation, drafting the article, approval of the article, and data collection.

## Notes

Conflict of Interest and Source of Funding: There are no conflicts of interest or funding sources to disclose.

### Competing Interest Statement

The authors have declared no competing interest.

